# The GEF-GAP pair Solo and DLC3 balances Rho activity during endocytic transport

**DOI:** 10.1101/2022.03.01.482527

**Authors:** Cristiana Lungu, Florian Meyer, Marcel Hörning, Jasmin Steudle, Anja Braun, Bettina Noll, David Benz, Felix Fränkle, Simone Schmid, Stephan A. Eisler, Monilola A. Olayioye

## Abstract

The control of intracellular membrane trafficking by Rho GTPases is central to cellular homeostasis. How defined pairs of guanine nucleotide exchange factors (GEFs) and GTPase-activating proteins (GAPs) come together to locally balance GTPase activation cycles in this process is nevertheless largely unclear. By performing a microscopy-based RNAi screen we here identify the RhoGEF protein Solo as a counterpart of DLC3, a RhoGAP protein with established roles in membrane trafficking. Biochemical, imaging and optogenetics assays further uncover Solo as a novel regulator of endosomal RhoB. Remarkably, we find that the interplay between Solo and DLC3 controls not only the activity but also total protein levels of RhoB. Together, the results of our study uncover the first functionally connected RhoGAP-RhoGEF pair at endomembranes and place the Solo-DLC3 axis at the core of endocytic trafficking.

## INTRODUCTION

Intracellular trafficking relies on a tight cooperation between cytoskeletal elements, proteins and lipids that come together to regulate cellular homeostasis (Anitei and Hoflack, 2011; Olayioye et al., 2019; Phuyal and Farhan, 2019). While the contribution of the small GTPases of the Rab and Arf families to membrane trafficking is well established, the involvement of Rho GTPases in this process is less understood. Since aberrant membrane trafficking is known to be associated with various diseases including cancer, it is important to elucidate the molecular mechanisms underlying the control of Rho GTPase signalling at different target membranes.

Rho GTPases function as molecular switches cycling between an active GTP-bound state and an inactive GDP-bound state. RhoGEFs promote the exchange of bound GDP for GTP, leading to Rho activation and initiation of downstream signalling pathways. RhoGAPs promote the low intrinsic GTP hydrolysis activity of the Rho GTPase, thereby dampening downstream signalling (Bos et al., 2007; Vigil et al., 2010). In addition to the plasma membrane, active Rho GTPase pools are also found at endomembranes such as endosomes and the Golgi complex (Olayioye et al., 2019; Phuyal and Farhan, 2019). For example, RhoB is detected at endosomes from where it controls the recycling of cargoes such as the epidermal growth factor receptor (EGFR) (Adamson et al., 1992; Phuyal and Farhan, 2019; Wherlock et al., 2004). Notably, RhoB pools localized at the plasma membrane and endosomes appear to be functionally distinct, the latter playing the main role in EGFR recycling (Wherlock et al., 2004). Additionally, RhoB and RhoA are important for the architecture of the Golgi complex, their hyperactivation causing severe fragmentation of this organelle (Zilberman et al., 2011). With 145 members, the RhoGEF and RhoGAP proteins greatly exceed the 10 classical Rho family GTPase switch proteins (Aspenström, 2020; Braun and Olayioye, 2015; Fritz and Pertz, 2016; Olayioye et al., 2019). This complexity underscores the need for unbiased approaches to identify and characterize defined pairs of GEFs and GAPs co-regulating Rho GTPase signalling in distinct cellular contexts.

Among the RhoGAP proteins, the Deleted in Liver Cancer (DLC) family stands out, being deregulated in different cancer types more frequently than any other Rho regulator (Kandpal, 2006; Xue et al., 2008). DLC3, also known as STARD8, is the least characterized member of the DLC family (Braun and Olayioye, 2015). Work from our lab has uncovered a RhoGAP-specific function of DLC3 in the maintenance of the integrity of the Golgi complex and the endocytic recycling compartment (Braun et al., 2015; Noll et al., 2019). Furthermore, in HeLa cells, DLC3 partially co-localized with endosomal RhoB and regulated trafficking in a RhoA/B dependent manner (Braun et al., 2015). The RhoGEF counterpart of DLC3 responsible for Rho activation at endomembranes has nevertheless remained elusive to date.

By performing an image-based RNAi screen using the morphology of the Golgi complex as a readout, we here identify the RhoGEF protein Solo, also known as ARHGEF40 or Scambio, as an antagonist of DLC3 activity. Subcellular fractionation, imaging and optogenetic recruitment experiments further revealed Solo to be present on a subset of RhoB-positive endosomes, where it can locally activate this GTPase. In addition to activity we unexpectedly find that Solo and DLC3 co-regulate RhoB protein levels. Together, the results of our study uncover the first functionally connected RhoGAP-RhoGEF pair at endomembranes and place the Solo-DLC3 axis at the core of RhoB regulation and endocytic trafficking.

## METHODS

### Cell culture

HEK293T and HeLa cells (ATCC) were cultured in RPMI 1640 (Invitrogen) supplemented with 10% fetal calf serum (FCS; PAA Laboratories, Cölbe, Germany). The cells were maintained in a humidified incubator at 37 °C with 5% CO_2_. All cell lines were authenticated, tested negative for Mycoplasma (Lonza, LT07-318) and were kept in culture for no longer than 2 months.

### Antibodies and reagents

Following primary antibodies were used for detection: anti-AKT (Cell Signalling Technology, Cat# 2920), anti-Phospho-(Thr308)-AKT (Cell Signalling Technology, Cat# 2965), anti-EEA1 (Cell Signalling Technology, Cat# 3288), anti-EGFR (Santa Cruz, Cat# sc-03-G), anti-EGFR (Santa Cruz, Cat# sc-101), anti-Phospho-(Tyr1068)-EGFR (Cell Signalling Technology, Cat# 3777), anti-p44/42 MAPK (Erk1/2) (Cell Signalling Technology, Cat# 9107), anti-Phospho- (Thr202/Tyr204) p44/42 MAPK (Erk1/2) (Cell Signalling Technology, Cat# 9101, anti-GAPDH (Sigma-Aldrich, Cat# G9545), anti-GFP (Cell Signalling Technology, Cat# 2956), anti-Giantin (Abcam, Cat# 37266), anti-GST (GE Healthcare, Cat# 27457701), anti-HA (Sigma-Aldrich, Cat# 11867423001), anti-Rab5 (Cell Signalling Technology, Cat# 3547), anti-Rab7 (Cell Signalling Technology, Cat# 9367), anti-RhoA (Santa Cruz, Cat# sc-418), anti-RhoB (Santa Cruz, Cat# sc-8048), anti-RhoB (Cell Signalling Technology, Cat# 2098), anti-RhoC (Cell Signalling Technology, Cat# 3430), anti-Solo (custom made, Pineda Antibody Service, Berlin).

HRP-labelled secondary anti-mouse-IgG, anti-rabbit-IgG and anti-goat-IgG were purchased from Dianova (Cat# 115-035-062, Cat# 111-035-144, Cat# 705-035-147). Alexa-Fluor^®^-labelled secondary IgG antibodies, Alexa-Fluor^®^-labelled phalloidin and TO-PRO-3 were obtained from Invitrogen. DAPI was obtained from Sigma-Aldrich.

### Generation of a polyclonal anti-Solo antibody by rabbit immunization

Protein epitope analyses including sequence comparisons were carried out to avoid cross-reactivity. The selected peptide (1015 – 1032aa of human ARHGEF40, isoform 1, UniProtKB Q8TER5-1) was synthesized and separated by HPLC for subsequent immunization of three rabbits (Pineda Antibody-Service, Berlin, Germany). The pre-immune and resulting immunization sera were analysed by western blotting, periodically over a total time span of three months, which included monthly rounds of immunization. To compare the quality, specificity and affinity of the sera collected from the three animals, lysates of HeLa cells with either Solo overexpression or Solo knockdown were used (see Figure S2B). The IgG fraction of the antibody generated was separated by affinity chromatography (Pineda Antibody-Service, Berlin, Germany) and aliquots were stored at −80°C for subsequent use.

### RNA interference and plasmid transfection

For transient knockdowns, the cells were transfected using Lipofectamine™ RNAiMAX (Invitrogen) according to manufacturer’s instructions. The cells were used for experiments at 48 – 72 h post transfection as described in the figure legends. The siRNAs used were: negative control siRNA [siNT, ON-TARGETplus^®^ non-targeting control pool D-001810-10 from Dharmacon (Lafayette, CO)], siDLC3sp (siGENOME SMARTpool human STARD8 M-010254 from Dharmacon), siDLC3ss (Silencer^®^ Select STARD8 s18826 from ThermoFisher Scientific), siARHGEF2sp (ON-TARGETplus^®^ SMARTpool, L-009883), siARHGEF2ss (Silencer^®^ Select ARHGEF2, 4392420 from ThermoFisher Scientific) siRhoBsp (ON-TARGETplus^®^ SMARTpool RhoB L-008395 from ThermoFisher Scientific), siSolo ss#2 (Silencer^®^ Select ARHGEF40 s31288 from ThermoFisher Scientific) and siSolo sp#4 (ON-TARGETplus Human ARHGEF40 siRNA J-030269-12 from Dharmacon). siRNAs sequences of the RhoGEF library are provided in Table S1_RhoGEF screen siRNAs seq. For plasmid transfections of HeLa and HEK293T cells, LipofectamineLTX(ThermoFisher Scientific) and TransIT-293 (Mirus) regents, respectively, were used according to manufacturer’s instructions.

### RNAi Golgi screen and analysis

The RhoGEF screen was performed using a WiScan Hermes High content imaging system (Idea Biomedical, Israel). To this end, 5 x 10^4^ HeLa cells were seeded per well in CollagenR coated glass bottom 96-well plates (Greiner, Germany). The cells were subjected to reverse transfection using 2.5 pmol siRNA consisting of siDLC3sp and siRhoGEF or siNT control mixed at a 1: 1 ratio. 72 h post-transfection, the cells were washed with PBS, fixed and stained for the Golgi complex (Giantin, Alexa Fluor^®^ 488, green channel) and the nuclei (TO-PRO-3, far red channel) as described in the immunofluorescence staining section. Images were acquired with the high resolution 40x 0.75 NA objective for green and red channels. Per well, a coverage of 85 % and a field density of 20 % was applied, resulting in 200 frames. Images were analysed with the implemented WiSoft Minerva analysis application development platform: In brief, nuclei and Golgi complexes were segmented using the corresponding channels. Cells were segmented related to nuclei and Golgi compartments using the cytoplasmic background signal of the Golgi staining. To increase the accuracy, cells touching the image edge, mitotic cells and cells with oversegmented nuclei were excluded from the analysis. The number and average perimeter of Golgi fragments were measured per cell in order to quantify the state of Golgi fragmentation. The screen was performed twice.

For statistical analysis, the number of Golgi vesicles and average perimeter of Golgi vesicles a log-transformation was applied, as the data show a Poisson distribution and strongly skewed normal distribution, respectively. The mean *μ*_log_ and standard deviation *σ*_log_ were calculated from the log-transformed data

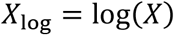

as

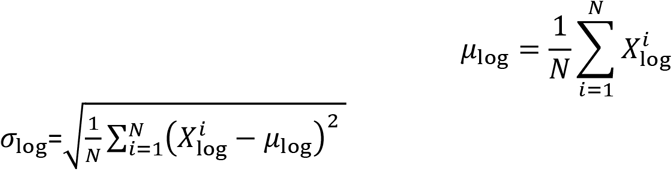

and back-transformed to obtain the mean and variance in normal space as

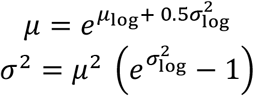

following the Finney estimator approach (Finney, 1941).

The data were statistically analysed using a N-way ANOVA (NANOVA) method in Matlab (Mathworks, R2021a). The two p-values obtained from the two screenings were combined as

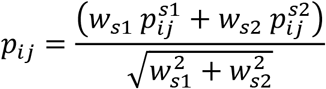

with the two weights

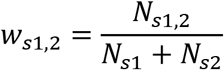

where *N* is the relative number of observables at each screening.

The Golgi screen dataset is available from the University of Stuttgart at: https://darus.uni-stuttgart.de/privateurl.xhtml?token=e0aa3f7b-ac8c-4253-8eb4-c8ae826346eb. The combined p values of the Golgi screen analysis are provided in Figure 1 – figure supplement 2B.

### DNA constructs

The GEF inactivating L1217E Solo point mutation was introduced by QuickChange site directed mutagenesis using pIRESNEO-Scambio-HA (Addgene plasmid #33354) as a template and the following forward and reverse primers: 5′- CTCGGTATGGGCGGGAGCTGGAGGAGCTCCTG -3′ and 5′- CAGGAGCTCCTCCAGCTCCCGCCCATACCGAG -3′. QuickChange site directed mutagenesis was also used to generate the RhoB Q63L (5′- GTGGGACACAGCTGGCCTGGAGGACTACGACCGC -3′ and 5′- GCGGTCGTAGTCCTCCAGGCCAGCTGTGTCCCAC -3′) and the RhoB T19N variant (5′- GATGGAGCCTGTGGAAAGAACTGCTTGCTCATAGTCTTC -3′ and 5′- GAAGACTATGAGCAAGCAGTTCTTTCCACAGGCTCCATC -3′) using the pEGFP-C1-RhoB plasmid as a template. The Opto-Endo Solo-GEF plasmid was generated by Gibson assembly whereby the coding sequence of PLD (Endosome-targeted optoPLD, (Tei and Baskin, 2020)) was exchanged for the Solo DHPH GEF domain (aa1076 – aa1380) amplified using the following primers : 5′- CGATCGGGCGGATCTATGGAGCGCAAGCGAAGC -3′ and 5′- GTTTGTAGCGCCGCTTCCCTCCTTGTTGTGGGCTGCC -3′. Solo GEF domain borders were selected following the approach previously used to generate the ARHGEF11 DHPH GEF domain (Valon et al., 2017). All constructs were validated by Sanger sequencing. The pCRV62-Met-Flag-DLC3 WT, mCherry-DLC3 K725E, pEGFP-C1-RhoB plasmids were previously described in (Braun et al., 2015; Holeiter et al., 2012; Noll et al., 2019). Endosome-targeted optoPLD was a gift from Jeremy Baskin (Addgene plasmid # 140056, (Tei and Baskin, 2020)). pEGFP-C1-Anillin AHPH WT and A70D/E758K (DM) were provided by Alpha Yap (University of Queensland, Australia) (Piekny and Glotzer, 2008; Priya et al., 2015). pcDNA™3.1 was from Invitrogen.

### Quantitative Real-Time PCR

Total RNA was isolated using the NucleoSpin^®^ RNA Kit (Macherey-Nagel, Hœrdt, France) according to manufacturer’s instructions. 100 ng RNA were used for real-time PCR with the Power SYBR^®^ Green RNA-to-C_T_™ 1-Step kit (Applied Biosystems) and the CFX96 Touch Real-Time PCR Detection System (Bio-RAD, 1855196). Changes in gene expression levels were determined relative to the housekeeping gene RPLP0 (5′- CTCTGCATTCTCGCTTCCTGGAG -3′ and 5′- CAGATGGATCAGCCAAGAAGG -3′) using the 2^−ΔΔCt^ method. Hs_ARHGEF40_1_SG and Hs_STARD8_1_SG QuantiTect Primer Assays (Qiagen) were used to detect the *ARHGEF40* and the *STARD8* transcripts, respectively. The 5′- CATTCTGACCACACTTGTACGC -3′ and 5′- GGTTTCTTTTCCCTCTCC TTGT -3′ primers were used for *RHOB*.

### Cell lysis, SDS-PAGE and western blotting

Cells were washed with PBS and harvested by scraping in ice-cold lysis buffer [150 mM Tris-HCl pH 7.5, 500 mM NaCl, 1% (v/v) Triton X-100, 0.5% (v/v) sodium deoxycholate, 0.1% (w/v) SDS, 10% Glycerol (v/v), 10 mM MgCl_2_, 1 mM DTT, 1 mM sodium orthovanadate, 10 mM sodium fluoride, 0.5 mM PMSF, 20 mM β-glycerophosphate and cOmplete™, EDTA-free Protease Inhibitor Cocktail (Roche)]. The cell lysates were passed 5 times through a 27-gauge needle, followed by 20 minutes incubation on ice. Subsequently, the whole-cell lysate was clarified by centrifugation for 10 min at 16,000 g and 4°C. The protein concentration of the supernatant was determined by performing a DC protein assay (Bio-rad). Equal protein amounts were separated by SDS-PAGE (NuPage^®^ Novex^®^ 4–12% Bis-Tris gels, Invitrogen) and transferred to nitrocellulose membranes using an iBlot^®^ device (iBlot^®^Gel Transfer Stacks; Invitrogen). The membranes were blocked with 0.5% (v/v) blocking reagent (Roche) in PBS containing 0.05% (v/v) Tween-20 and 0.01% (v/v) Thimerosal and then incubated with primary antibodies, overnight at 4°C, followed by 1 h incubation with HRP-conjugated secondary antibodies at room temperature. The chemiluminescence signal was detected using an AmershamTM Imager 600 device (GE Healthcare) followed by quantification of the 16-bit images in the linear range using the inbuilt ImageQuant TL 8.1 software.

For inhibitor experiments, HeLa cells were treated with either 30 mM NH_4_Cl (Carl Roth) or 5 μM MG132 (Selleck Chemicals) for 12 h before harvesting. Cycloheximide (Calbiochem) was used for the indicated time points at a concentration of 50 μg/mL.

### RBD pulldowns

BL21 – CodonPlus competent cells (Agilent Technologies) were transformed with a plasmid encoding the GST-tagged domain of Rhotekin (Addgene plasmid # 15247). Cells were cultivated in Terrific broth (TB) medium at 37°C while shaking until an OD (600 nm) of 0.6–0.7 was reached. Then, protein expression was induced by addition of 1 mM of isopropyl-β-D-thiogalactoside and the culture was incubated at 21°C shaking at 200 rpm for another 16 h. The cells were harvested by centrifugation at 4°C (15 min at 4600 rpm) and the pellet resuspended in sonication buffer [50 mM Tris-HCl pH 7.5, 500 mM NaCl, 1 mM DTT, 5 mM MgCl_2_, 10% glycerol, cOmplete™, EDTA-free Protease Inhibitor Cocktail (Roche)]. The cells were lysed by sonication and centrifuged at 18 000 rpm for 1 h to prepare a clear lysate, which was applied on a GST-sepharose column (GE Healthcare). After washing with sonication buffer, the protein was eluted with sonication buffer containing 50 mM reduced glutathione (Sigma-Aldrich) and dialyzed against storage buffer [50 mM Tris-HCl pH 7.5, 150 mM NaCl, 1 mM DTT, 5 mM MgCl_2_, 10% glycerol] for 3 h. The eluted protein was aliquoted and stored at −80°C till further use.

For pulldowns, cells were stimulated with 100 ng/ml EGF for 5 min, washed once with cold TBS [50 mM Tris-HCl pH 7.6, 140 mM NaCl] and lysed in ice-cold RBD lysis buffer [150 mM Tris-HCl pH 7.5, 500 mM NaCl, 1% (v/v) Triton X-100, 0.1% (w/v) SDS, 0.5% sodium deoxycholate (v/v), 1mM DTT, 10% Glycerol (v/v), 10mM MgCl_2_, 1 mM EGTA, 1 mM sodium orthovanadate, 10 mM sodium fluoride, 0.5 mM PMSF, 20 mM β-glycerophosphate and cOmplete™, EDTA-free Protease Inhibitor Cocktail (Roche)]. The cell lysate was passed 5 times through a 27-gauge needle, followed by centrifugation for 5 min at 16,000 g and 4°C. Equal amounts of cleared lysates were incubated for 45 min at 4°C with 30 μg GST–RBD protein pre-bound to GST-sepharose beads. Beads were rapidly washed four times with RBD lysis buffer, the eluted proteins were separated by SDS-PAGE and analysed by western blotting as described above.

### Immunofluorescence staining and confocal microscopy

Cells grown on glass coverslips coated with 10 μg/ml collagen R (Serva; Heidelberg, Germany) were fixed for 15 min with 4% (w/v) paraformaldehyde. After washing with PBS supplemented with Ca^2+^ and Mg^2+^, the cells were incubated for 15 min with 150 mM glycine in PBS followed by permeabilization for 5 min with 0.1% (v/v) Triton X-100 in PBS. Blocking was performed with 5% (v/v) goat serum (Invitrogen) in PBS containing 0.1% (v/v) Tween-20. Fixed cells were incubated with primary antibodies diluted in blocking buffer overnight at 4°C. Following three washing steps with PBS, the cells were then incubated with Alexa-Fluor^®^-(488, 546 or 633)- labelled secondary antibodies in blocking buffer for 1 h at room temperature. For the immunolabelling of endogenous RhoB, the method by (Adamson et al., 1992) was followed. In short, after fixation and permeabilization, cells were incubated with 10% FCS in PBS for 30 min, followed by incubation overnight at 4°C with the anti-RhoB antibody (Santa Cruz, Cat# sc-8048) diluted 1:100 in incubation buffer (50 mM Tris-HCl, pH 7.6, 1% NP-40, 0.5% sodium deoxycholate and 5 mM EDTA), which was also used for washing and dilution of the secondary antibody. Nuclei were counterstained with DAPI and the coverslips were mounted in Molecular Probes™ ProLong™ Gold Antifade mountant (ThermoFisher Scientific).

All samples were analysed at room temperature using a confocal laser scanning microscope (LSM 710, Carl Zeiss; Oberkochen, Germany) equipped with a Plan Apochromat 63×/1.40 DIC M27 (Carl Zeiss) oil-immersion objective. The excitation wavelengths and detection window used were as follows: 405nm and 425nm-488nm for DAPI; 488nm and 496nm-544nm for GFP and Alexa-Fluor^®^-488 coupled probes; 561nm and 573nm-621nm for mCherry and Alexa-Fluor^®^-546 coupled probes; 633nm and 650nm-717nm for Alexa-Fluor^®^-633 coupled probes. All images were acquired with the pinhole of each channel adjusted to the same optical slice thickness. Linear adjustments to brightness, contrast and maximum intensity projections were made using the ZEN software (Carl Zeiss). Fluorescence intensities along a line of interest were measured using the ZEN software.

### Setup of optogenetics experiments used to validate the OptoEndo-Solo-GEF tool

Samples were illuminated with 405 nm light in a custom-made box equipped with 6 equally spaced high-power 3W LEDs 700mA. Illumination cycles were 5 sec on – 35 sec off. Following 90 cycles of illumination, cells were fixed with 4% (w/v) paraformaldehyde. This was followed by permeabilization for 5 min with 0.1% (v/v) Saponin in PBS before immunofluorescence staining for EEA1, as described above. Dark control samples were not exposed to light and were fixed directly after being taken out of the incubator.

### Setup and analysis of live cell imaging optogenetics experiments

HeLa cells were seeded onto collagen-coated 35 mm high glass bottom μ-Dishes (ibidi, cat. no: 81158). After 48 h, cells were transfected with 1.75 μg total plasmid DNA mixture consisting of 0.6 μg OptoEndo-Solo-GEF, 0.25 μg pEGFPC1-AHPH and 0.65 μg pcDNA3. The cells were imaged 6 h post transfection. For knockdown experiments, the cells were reverse transfected with RhoB siRNA 48 – 72 h before transfection with plasmid DNA. Images were acquired every 20 s on an LSM910 Airyscan (Carl Zeiss) instrument equipped with a Plan-Apochromat 63x/1.4 Oil DIC objective and an environment control chamber. The excitation wavelengths and detection window used were 488 nm and 499-549 nm for GFP and 561 nm and 573-627 nm for mCherry, respectively. For imaging, only cells with weak basal mCherry fluorescence were selected. While identifying pEGFPC1-AHPH double transfected cells, the exposure time to the 488 nm laser was kept to a minimum. Several imaging dishes were prepared in parallel for each condition to avoid unintentional optogenetic recruitment of OptoEndo-Solo-GEF outside of the acquisition timeframes.

To identify and follow single endosomes, the plugin TrackMate (Tinevez et al., 2017) was employed. To this end, .czi files were imported into Fiji using Bio-Formats (Linkert et al., 2010) and the LoG detector was used to detect individual endosomes in the pre-processed (median) mCherry channel. The expected vesicle diameter was set to 0.7 μm and the quality threshold to 50. To ensure that the detected endosomes are linked correctly during tracking, the linking max distance was set to 1.5 μm and the gap closing max distance/max frame gap to 0. Only endosomes detected over 20 consecutive time frames, including the first acquisition frame, were taken into consideration for analysis.

To define endosomes as positive for the GFP-AHPH sensor, the maximum GFP intensity was used. This was because the mCherry and the GFP signal did not show complete overlap, which could bias analysis based on mean fluorescence intensity. To calculate the median background GFP signal for normalization, the maximum GFP intensity of all endosomal ROIs per cell was extracted. During tracking, each maximum GFP intensity per frame was used to calculate the mean maximum GFP intensity per track. All calculations were done using an in house written python script.

### Live cell imaging

HeLa cells were seeded onto collagen-coated 35 mm high glass bottom μ-Dishes (ibidi, cat. no: 81158) followed by transient transfection with the plasmids of interest as described in the RNA interference and transfection section above. 24 h later the cells were imaged on a Zeiss Cell Observer SD Spinning Disk microscope equipped with an EMCCD camera (Photometrics Evolve 512). GFP was visualized with a 488 nm diode laser in combination with a FE01-520/35 nm emission filter. mCherry was visualized with a 561 nm diode laser in combination with a 600_50 ET emission filter. Images were acquired at 37°C and 5% CO_2_ every 20 s for a time interval of 10 min. Image processing was done with Zen 2.3 blue software.

### EGFR trafficking and signalling assays

For immunofluorescence analysis of EGFR trafficking, HeLa cells were starved for 20 h in RPMI supplemented with 0.5% serum before stimulation with 10 ng/mL EGF. The cells were then washed with PBS, fixed, blocked in 5% (v/v) goat serum in PBS and incubated with an antibody recognising the extracellular domain of EGFR (Santa Cruz, Cat# sc-101). This was followed by permeabilization as described above and incubation with an antibody recognising the C-terminus of EGFR (Santa Cruz, Cat# sc-03-G). Cells used for EGFR signalling analysis were treated in a similar manner before being lysed in ice-cold RIPA buffer [50 mM Tris-HCl pH 7.5, 150 mM NaCl, 1% (v/v) NP-40, 0.1% (w/v) SDS, 0.25% sodium deoxycholate (v/v), 1 mM EDTA, 1 mM sodium orthovanadate, 10 mM sodium fluoride, 0.5 mM PMSF, 20 mM β-glycerophosphate and cOmplete™, EDTA-free Protease Inhibitor Cocktail (Roche)] and processed for western blot analysis.

### Statistical analysis

Data are presented as mean + SEM, where each experiment, apart from the Golgi screen, was performed at least 3 times. The Golgi screen was performed twice. For box plots, the box represents the 25–75th percentiles, and the median is indicated; whiskers extend 1.5 times the interquartile range from the 25th and 75th percentiles, and outliers are represented as determined by GraphPad Prism 7 software. ‘N’ refers to the number of sample points and ‘n’ to the number of independent experiments. Significance between multiple groups was determined by one-way ANOVA followed by Tukey (all vs all) or Dunnett (all vs control) multiple comparison post-test, as detailed in the figure legends. The two-way ANOVA analysis was followed by Sidak post-testing. Significance between two groups was determined by t-test. Data were analysed using GraphPad Prism 7. p values: p ≤0.03 (*), p ≤0.002 (**), p ≤ 0.0002 (***), p ≤ 0.0001 (****), n.s.: non-significant. See also Table S2_Summary p values.

## RESULTS

### RhoGEF screen identifies Solo as a potential DLC3 antagonist

We previously found that DLC3 controls the integrity of the Golgi and endocytic recycling compartments in a RhoA/B dependent manner (Braun et al., 2015). To identify antagonists of DLC3 function, we screened twenty-three GEF proteins with known in vitro, and in many cases, also in vivo specificity for the RhoA/B/C subfamily of GTPases using the morphology of the Golgi complex as a readout. To this end, HeLa cells were transiently transfected with Smartpool siRNAs specific for the individual RhoGEFs along with DLC3-specific siRNA, followed by automated image analysis of immunostained cells. Labelling of the Golgi and the nucleus were used to identify and segment the cells of interest as well as to detect outliers (Figure 1 – figure supplement 1-A and B). The morphological features of the Golgi complex, e.g. the number and average perimeter of Golgi fragments (Figures 1A and 1B) were then computed. In accordance with our previous work (Braun et al., 2015), DLC3 knockdown cells displayed a doubling of the number of giantin-positive fragments per cell and a significant decrease in the average perimeter of the Golgi fragments (Figures 1B, 1C and Figure 1 – figure supplement 2B). Remarkably, whereas the knockdown of most of the RhoGEF proteins had little effect on the number and the average perimeter of Golgi fragments in DLC3-depleted cells, the knockdown of the RhoGEF Solo stood out for its ability to bring the compact structure of the Golgi complex closest to the control cells (Figures 1A, 1B, and Figure 1 – figure supplement 1C and Figure 1 – figure supplement 2). This rescue was not due to a decrease in DLC3 knockdown efficiency upon simultaneous depletion of Solo (Figure 1 – figure supplement 1D).

**Figure 1:**
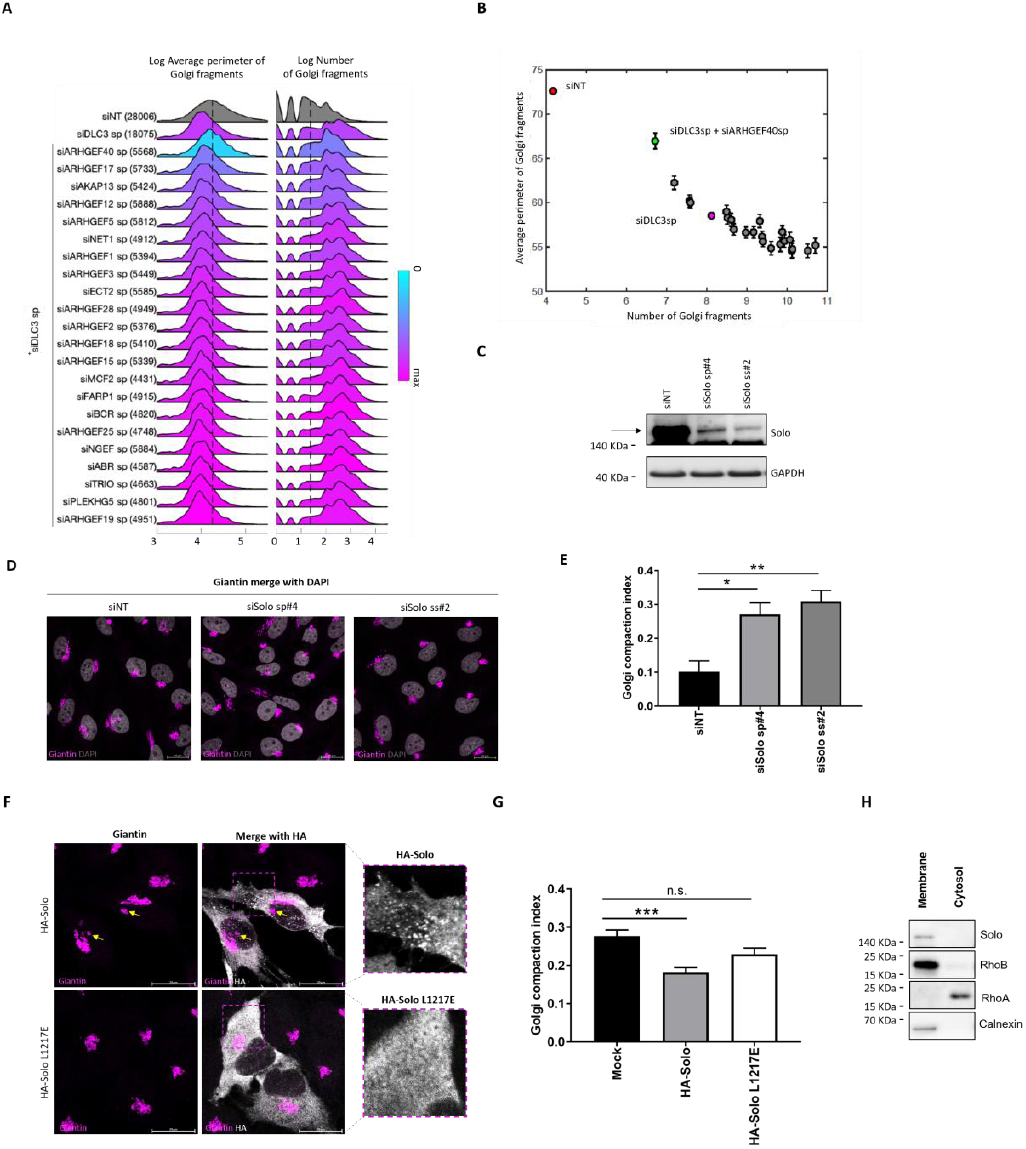
Imaging-based Golgi screen identifies the RhoGEF Solo as a potential DLC3 antagonist. (A-B) HeLa cells were transfected with control (siNT), DLC3-specific, or a combination of DLC3 and RhoGEF specific smartpool siRNAs. 72 h post transfection, cells were fixed, stained, and scanned with the Hermes WiScan. Images were analysed with WiSoft. (**A**) Quantification of the morphology of the Golgi complex for the RhoGEF screen performed as described above. Shown are the average perimeter of Golgi vesicles (left panel) and the number of Golgi fragments (right panel). The number of cells used for analysis is annotated in parenthesis for each siRNA transfection set. The colour scheme indicates the differences between the means of the control (siNT) and the respective knockdown. Statistical parameters, e.g. mean and variance, were obtained by log-transformation using the Finney estimator approach. (**B**) The relation between the number and the average perimeter of Golgi vesicles: Error bars = 95% confidence intervals. Data points highlighted in colour are the control (siNT, red), siDLC3sp (pink), and siDLC3sp + siARHGEF40sp (green) cells. Data were statistically analysed using the N-way ANOVA (NANOVA) method in Matlab. See also the 2D histograms of the number and average perimeter of each respective siRNA transfection (Figure S2A) and the p-value matrix for the specific statistical comparisons performed for each screen (Figure S2B). (**C**) Validation of the siRNAs used for Solo knockdown. HeLa cells were transfected with either control or two independent Solo siRNAs, followed by harvesting at 72 h post transfection. Cell lysates were analysed by immunoblotting with a Solo specific antibody. The band corresponding to Solo is annotated with an arrow. GAPDH was used as loading control. (**D**) Representative confocal immunofluorescence microscopy images showing the morphology of the Golgi apparatus (Giantin, pink) in HeLa cells 72h post transfection with either control or Solo specific siRNAs. Nuclei were stained with DAPI (grey). All images were acquired and displayed using identical settings. (**E**) Quantification of the morphology of the Golgi complex in HeLa cells exemplarily shown in (D) using the Golgi compaction index method (Bard et al., 2003). Data show the mean + S.E.M. (n = 3, N≥30 cells). Statistical comparison by one-way ANOVA and Dunnett post-test; *p≤ 0.03, **p≤ 0.002. (**F**) Representative confocal immunofluorescence microscopy images documenting the morphology of the Golgi apparatus (Giantin, pink) in HeLa cells 24 h post transient transfection with plasmids encoding either wild type or GEF inactive L1217E HA-Solo (HA, white). Note the fragmentation of the Golgi complex observed selectively in cells expressing wild type Solo (yellow arrows). All images were acquired and displayed using identical settings. (**G**) Quantification of the Golgi morphology in HeLa cells with transient overexpression of wild type and L1217E HA-Solo as shown in (F). The box plot shows the results of three independent experiments. Centre lines show the medians; box limits indicate the 25th and 75th percentiles as determined by GraphPad Prism 7 software; whiskers extend 1.5 times the interquartile range from the 25th and 75th percentiles, outliers are represented by dots. N = 37 – 46. The significance of differences was analysed by a one-way ANOVA and Dunnett post-test; ***p≤ 0.0002, n.s.: non-significant. Scale bars: 20 μm. (**H**) HeLa cells were homogenised and fractionated into the membrane and cytosol fractions. Samples were analysed by immunoblotting using the indicated antibodies. See also Figures 1 – supplement 1 – 3 and Tables S1-S3.

### Solo is required for the integrity of the Golgi complex in a RhoGEF-dependent manner

To validate Solo as a regulator of Golgi complex morphology, we depleted the transcript using two independent siRNAs and confirmed the efficient knockdown at the RNA (Figure 1 – figure supplement 3A) and protein level using a Solo-specific antibody generated in-house (Figure 1C and Figure 1 – figure supplement 3B). For both siRNAs, Solo knockdown led to a significant increase in the Golgi compaction index, without affecting total cell area (Figures 1D, 1E, Figure 1 – figure supplement 1E and F). To investigate if Golgi morphology changes are direct and depend on Solo GEF activity, we transfected HeLa cells with plasmids encoding either the wild-type protein or a GEF inactive Solo mutant (Abiko et al., 2015). Only the expression of wild-type Solo led to a significant fragmentation of the Golgi complex (Figures 1F and 1G). These findings consolidate the results of the screen and show that Solo plays a previously unappreciated role in maintaining the integrity of the Golgi compartment.

We further observed that wild-type Solo displayed a vesicular localisation while the GEF inactive protein was distributed diffusely throughout the cytoplasm (Figure 1F, crop-ins). Since the Solo antibody did not show a specific signal in immunostainings of HeLa cells, we employed subcellular fractionation to monitor the distribution of endogenous Solo in the cytosolic vs. membrane fraction, containing e.g. endosomes and the plasma membrane (Jia and Bonifacino, 2019) (Figure 1H). Calnexin, which was used as a marker for the membrane fraction showed the expected extraction pattern and validated our approach. Interestingly, endogenous Solo was predominantly detected in the membrane fraction together with the small GTPase RhoB but not RhoA. These results support the involvement of Solo in membrane trafficking and hint at a role of this RhoGEF in RhoB regulation.

### Solo activates RhoB on endosomes

A characteristic of small GTPases is to bind to their GEFs in a GDP-bound or nucleotide-free state. By contrast, effectors bind stronger to the GTP-loaded form. Since Solo has not been previously linked to RhoB, we performed GFP pulldown assays using lysates from HEK293T cell expressing HA-Solo and conformationally locked RhoB-GFP variants (Figure 2A). As anticipated based on its function as a RhoGEF, Solo associated predominantly with the RhoB 19N GDP-locked mutant and less with wild-type RhoB or Q63L RhoB, mimicking the constitutively active protein. No signal was detected for the GFP control, validating the specificity of the interaction. To gain spatial insights into the Solo – RhoB association we next performed immunofluorescence imaging of HeLa cells transiently expressing HA-Solo and GFP-RhoB 19N. In line with previous reports (Gong et al., 2018), GDP-locked RhoB localised on vesicles known to be endosomes (Figure 2B). Notably, Solo was detected on a subset of the RhoB 19N positive vesicles. Together with the pulldown results, this finding supports the hypothesis that Solo regulates RhoB activity on endosomes.

**Figure 2:**
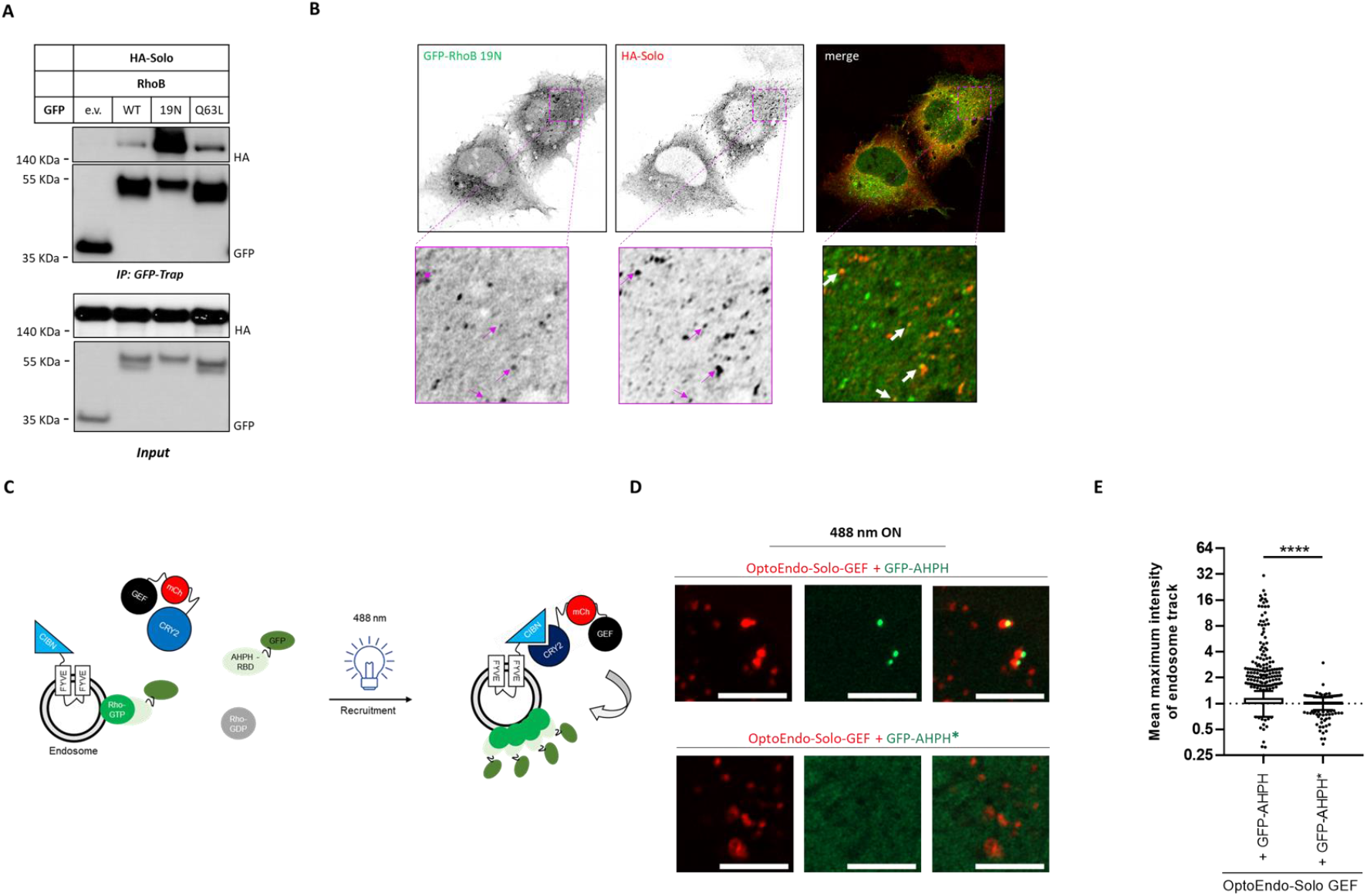
Solo activates RhoB on endosomes. (**A**) HEK293T cells were co-transfected with plasmids encoding for HA-Solo and the indicated GFP-RhoB variants. A plasmid encoding for empty GFP served as a negative control. 24 h post transfection the lysates were analysed directly (input) or used for GFP-Trap assays followed by immunoblotting with the indicated antibodies. (**B**) Representative confocal immunofluorescence microscopy images of HeLa cells transiently expressing GFP-RhoB 19N (green) and HA-Solo (red). Arrows denote exemplary areas of co-localisation between the two proteins. Images show one confocal plane. (**C**) Schematic representation of the optogenetic experimental workflow employed to monitor endosomal RhoB activation upon Solo-GEF recruitment. See text for details. (**D**) 6 h post transfection with the constructs described in (C), HeLa cells were exposed to 488-nm light and imaged. Shown are representative confocal microscopy images, illustrating the enrichment of the GFP-AHPH Rho-GTP reader (green) to sites of OptoEndo-Solo-GEF (red) recruitment. The GFP-AHPH A70D/E758K mutant (GFP-AHPH*) was used as a negative control. All images were acquired and are displayed with the same settings. Images show one confocal plane. Scale bar: 5 μm. (**E**) Quantification of the GFP-AHPH recruitment experiment shown in (D). The box plot shows the results of three independent experiments, where each dot represents one mCherry positive endosome. Centre lines show the medians; box limits indicate the 25th and 75th percentiles as determined by GraphPad Prism 7 software; whiskers extend 1.5 times the interquartile range from the 25th and 75th percentiles, outliers are represented by dots. N = 30 cells; n =3, with more than 600 vesicles analysed per condition. The significance of differences was analysed by Mann-Whitney t-test; ***p≤ 0.0002, n.s.: non-significant. See also Figure 2 – figure supplement 1. For statistical testing data see Table S2.

To address this hypothesis, we generated OptoEndo-Solo-GEF as a tool to assess the direct activation of endosomal RhoB upon targeted recruitment of the Solo GEF domain. Specifically, in this optogenetic approach, CRY2 was fused to the Solo DHPH-GEF domain and mCherry, while CIBN was fused to the 2xFYVE domain of Hrs1 (Tei and Baskin, 2020) (Figure 2C). The latter recognizes PtdIns(3)P, which is present on the membranes of early and late endosomes (Raiborg et al., 2013). In the absence of blue light, CRY2-Solo-GEF-mCherry displayed a weak mCherry fluorescence, being localized diffusely through the cytoplasm (Figure 2 – figure supplement 1A). By contrast, acute illumination resulted in the rapid recruitment of the probe to vesicles, which partially co-localised with the early endosomal marker EEA1 (Figure 2 – figure supplement 1A and 1B). To specifically detect Rho-GTP, we co-expressed the GFP-AHPH biosensor (Piekny and Glotzer, 2008; Priya et al., 2015). Strikingly, OptoEndo-Solo-GEF recruitment resulted in a significant local enrichment of the GFP-AHPH sensor on endosomes but not of the GFP-AHPH A70D/E758K binding pocket mutant (Figures 2D and E). While the GFP-AHPH probe does not distinguish between RhoA, RhoB and RhoC, RhoB is the only member that localises to endosomes (Fernandez-Borja et al., 2005). Our findings thus provide support for the GEF activity of Solo regulating RhoB at endosomal membranes.

### Solo and DLC3 co-regulate RhoB activity and protein levels

We next investigated how the interplay between Solo and DLC3 converges on RhoB by first performing live cell imaging experiments in HeLa cells transiently co-expressing fluorescently tagged Solo and DLC3. To preserve cell morphology, the K725E GAP-inactive DLC3 mutant, which recapitulates the localisation of wild-type DLC3, was used (Holeiter et al., 2012). Spinning disk confocal microscopy revealed dynamic co-localisation events that occurred between GFP-Solo and mCherry-DLC3 K725E in particular on vesicular-like structures at cell protrusions (Figures 3A, 3B and Figure 3 – figure supplement 1). Unfortunately, while both Solo and DLC3 individually co-localised with RhoB in HeLa cells (Figure 2B) (Braun et al., 2015), we were not able to detect their co-localization in triple transfection experiments. This is probably due to the rapid turnover rate of the GEF – GAP - Rho complex.

**Figure 3:**
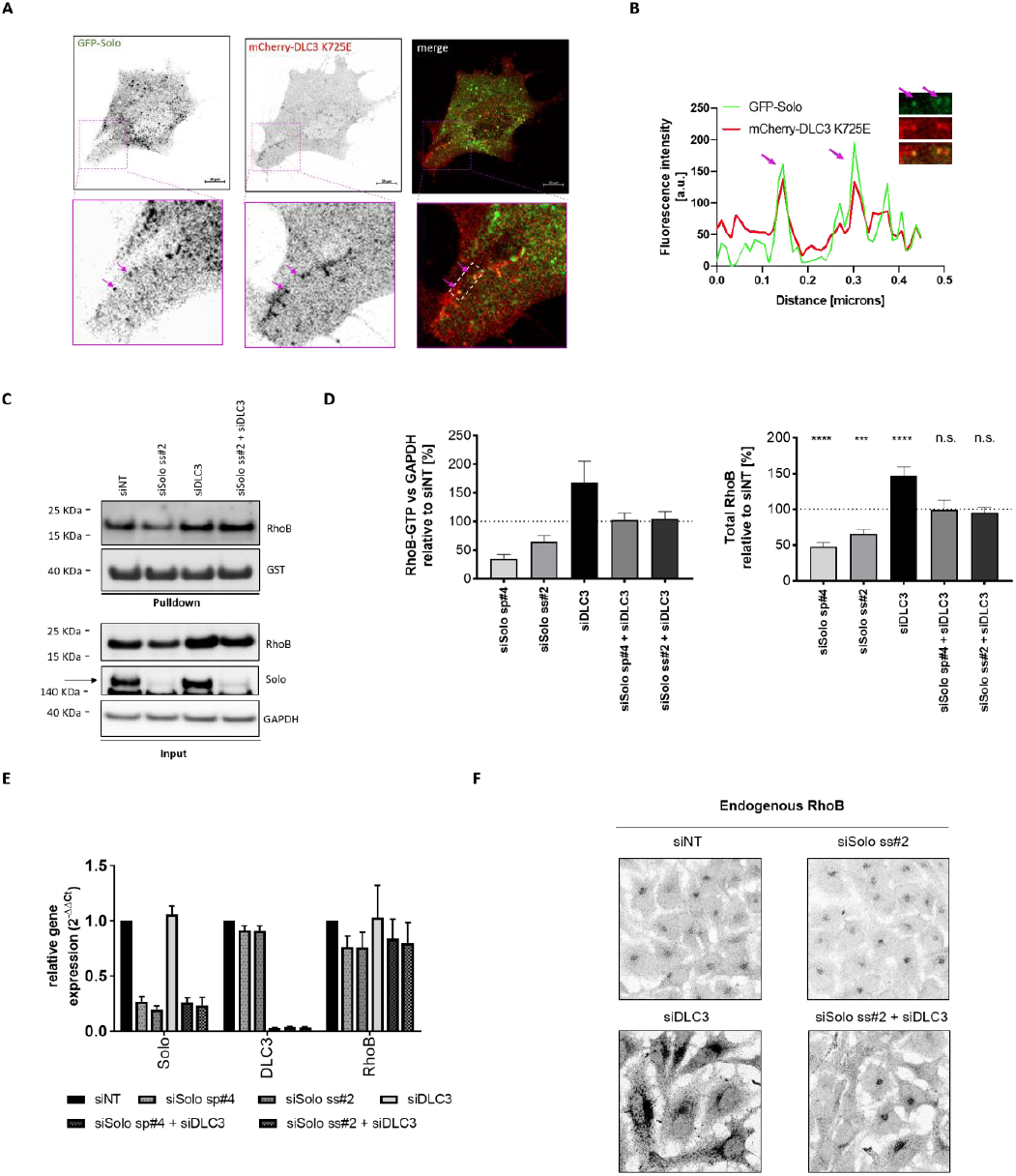
Solo and DLC3 co-regulate RhoB activity and protein levels. **(A)** HeLa cells were transiently transfected with plasmids encoding GFP-Solo and mCherry-DLC3 K725E followed by imaging 24 h later. Shown is a selected time frame from the live cell imaging experiment. Arrows highlight exemplary areas of colocalisation between the two fluorescently tagged proteins. Images display a single confocal section. Scale bar: 10 μm. See also Movie S1. (**B**) Intensity profiles of GFP-Solo (green) and mCherry-DLC3 K725E (red) extracted in ImageJ along a straight line crossing the white rectangle annotated in (A). Arrows highlight exemplary areas of colocalisation between the two fluorescently tagged proteins. (**C**) HeLa cells were transfected with the indicated siRNAs. At 72 h post knockdown, the cells were stimulated with 100 ng/ml EGF for 5 min, followed by lysis and GST–RBD pulldown assays. Pulldowns and total cell lysates were analysed by immunoblotting with the indicated antibodies. The band corresponding to Solo is annotated with an arrow. (**D**) Western blots from the experimental set representatively shown in C were quantified using the ImageQuant TL software. (left) The RhoB-GTP pulldown signal was normalized to GAPDH and further divided by the signal obtained for the siNT sample. The value obtained for the siNT control was set as 100%, marked with the dotted line. (right) The total RhoB signal in the input was normalised to GAPDH and further divided by the signal obtained for the siNT sample. The value obtained for the siNT control was normalised to 100%, marked with the dotted line. The significance of differences was analysed by a one-way ANOVA and Dunett post-test; ****p≤ 0.0001, **p≤ 0.002, n.s.: non-significant. Control is the siNT sample. n = 3 – 5. (**E**) qRT-PCR measurements of the Solo, DLC3 and RhoB transcripts in HeLa cells subjected to knockdown with the indicated siRNAs. RNA was isolated at 72 h post transient transfection. Gene expression values were normalized to the reference gene RPLP0. Shown is the relative change of each transcript normalized to the corresponding siNT sample, which was set to 1. n = 3, error bars: SEM. (**F**) HeLa cells were transfected with the indicated siRNAs. At 72 h post knockdown, the cells were fixed and stained for RhoB. Shown are representative maximum intensity projections of 5 confocal sections. All images were acquired and displayed using identical settings. See also Figure 3 – figure supplements 1-3. For statistical testing data see Table S2.

We then used RBD pulldowns to directly assess how endogenous RhoB activity is regulated by Solo and DLC3. These assays revealed a 40% reduction in the total cellular GTP-RhoB levels (normalized to GAPDH) upon Solo depletion indicating that Solo activates endogenous RhoB in HeLa cells. Intriguingly, this decrease in RhoB-GTP was paralleled by a significant drop in RhoB protein levels (Figures 3C and 3D), which was not observed for RhoA or RhoC (Figure 3 – figure supplement 2A). We also measured Solo protein levels in HeLa cells depleted of GEFH1, one of the few established GEFs for RhoB (Kamon et al., 2006). No changes in RhoB amounts were observed under these conditions, indicating that the downregulation of RhoB is specific to Solo and not a general response to global perturbations of RhoB GEF activity (Figure 3 – figure supplement 2D). Importantly, simultaneous knockdown of Solo and DLC3 rescued RhoB-GTP as well as total RhoB amounts to levels that were comparable to those of control cells (Figures 3C and 3D). While normalisation of RhoB-GTP to RhoB-GDP amounts showed no net changes in RhoB activity among the different knockdowns (Figure 3 – figure supplement 2C), these results clearly demonstrate that Solo and DLC3 co-regulate RhoB in HeLa cells. These effects were predominantly observed at the post-transcriptional level as qPCR analysis did not reveal significant changes in RhoB transcript levels in response to neither Solo nor DLC3 depletion (Figure 3E).

An intact endosomal trafficking system was found to be required for proper RhoB subcellular localisation and the regulation of RhoB stability (Gong et al., 2018; Zaoui et al., 2019). When visualising endogenous RhoB using a specific RhoB antibody (Figure 3 – figure supplement 2C), we noted that RhoB was concentrated in the perinuclear area in control cells, while the stabilised RhoB protein also accumulated at the periphery in DLC3 knockdown cells (Figure 4F). Importantly, simultaneous knockdown of Solo not only reduced the increase in RhoB levels observed upon DLC3 depletion, but also led to re-clustering of the GTPase in the perinuclear area as observed for the control cells. This result demonstrates the Solo and DLC3 are required for maintaining the subcellular distribution patterns of RhoB.

**Figure 4:**
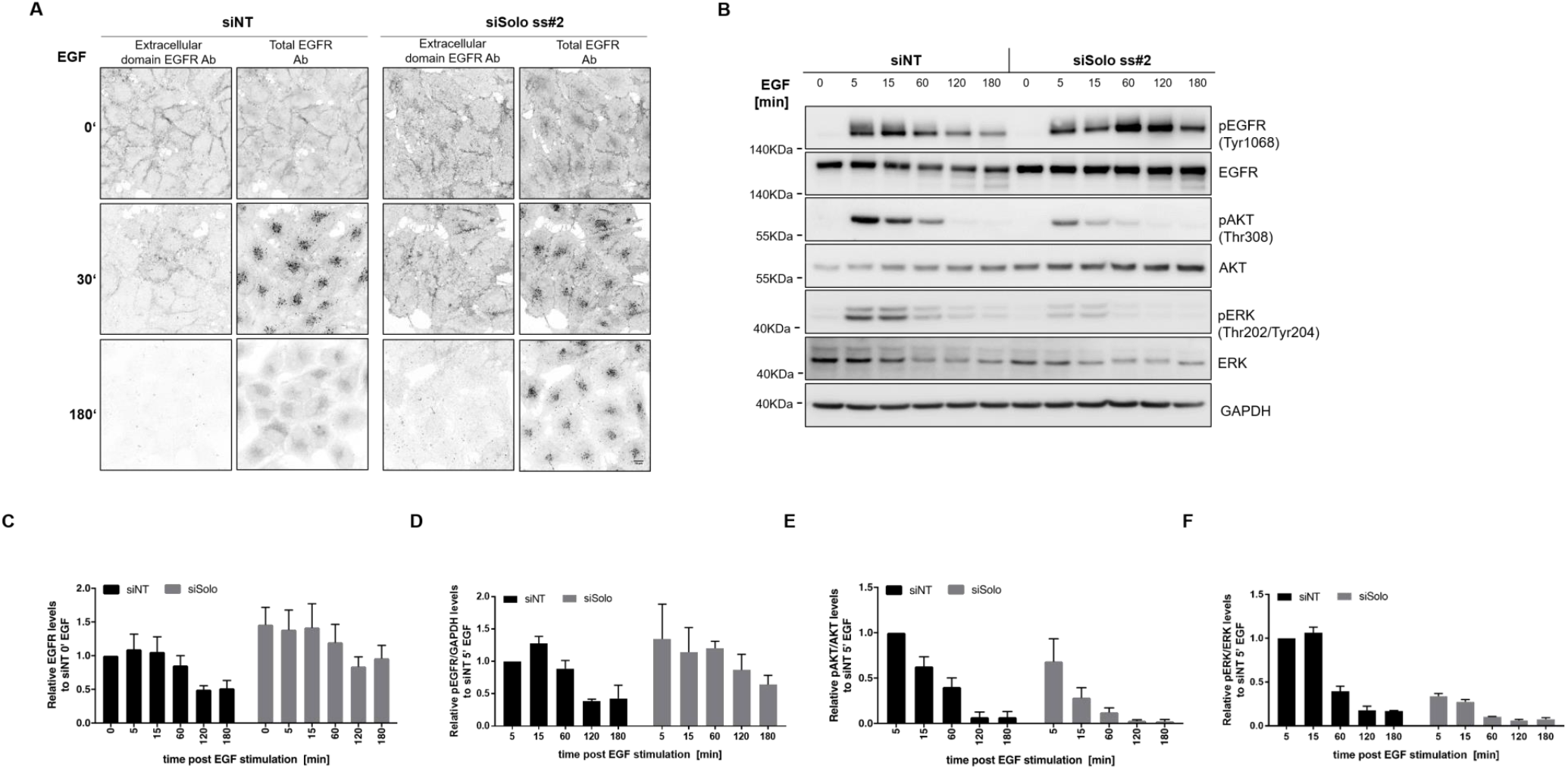
Solo knockdown delays EGFR trafficking and dampens the EGF signalling response. (**A**) HeLa cells were transiently transfected with control (siNT) or Solo-specific (siSolo ss#2) siRNAs. Following a 48 h incubation period, the cells were serum starved overnight and, prior to fixation, stimulated with 10 ng/ml EGF for the indicated times. The cells were then stained with an antibody recognising the extracellular domain of EGFR (surface EGFR). This was followed by permeabilization and immunostaining with an antibody recognising the C-terminal part of EGFR (total EGFR). Shown are representative maximum intensity projections of 5 confocal sections. All images were acquired and displayed using identical settings. Scale bar: 10 μm. (**B**) Following knockdown and EGF stimulation as described in A, HeLa cells were lysed, followed by immunoblotting with the indicated antibodies. The shown experiment is representative of three independent biological repeats. (**C**) Densitometric quantification of total EGFR levels in siNT and siSolo cells of the EGF stimulation experiment representatively shown in B. EGFR levels of the different time points were normalised to GAPDH and subsequently divided by the signal measured for the unstimulated siNT cells, which was set to 1. (**D**) Densitometric quantification of pEGFR levels in siNT and siSolo cells of the EGF stimulation experiment representatively shown in B. pEGFR levels of the different time points were normalised to GAPDH and subsequently divided by the signal measured for the siNT cells at 5 min post EGF stimulation, which was set to 1. (**E**) Densitometric quantification of the pAKT-Thr308 levels in siNT and siSolo cells of the EGF stimulation experiment representatively shown in B. The phosphorylation signal was normalised to the one obtained for the unphosphorylated protein and GAPDH and subsequently divided by the signal measured for the siNT cells at 5 min post EGF stimulation. The value obtained for the siNT cells was further normalised to 1. (**F**) Densitometric quantification of the pERK (Thr202/Tyr204) levels in siNT and siSolo cells of the EGF stimulation experiment representatively shown in B. The phosphorylation signal was normalised to the one obtained for the unphosphorylated protein and GAPDH and subsequently divided by the signal measured for the siNT cells at 5 min post EGF stimulation. The value obtained for the siNT cells was further normalised to 1. (C-F) The bar diagrams display the mean values of three independent biological repeats, cumulating two experiments using siSolo ss#2 and one experiment using siSolo sp#4. Error bars: SEM. See also Figure 4 – figure supplement 1. For statistical testing data see Table S2.

To obtain mechanistic insights into the regulation of RhoB protein levels, we next performed cycloheximide chase experiments (Figure 3 – figure supplement 3A and 3B). In agreement with measurements in other cell lines (Engel et al., 1998; Lebowitz et al., 1995), RhoB was found to be short-lived in HeLa cells, with a half-life of 2 ± 0.28h. By contrast, cells with DLC3 knockdown displayed higher RhoB protein stability, with the half-life increased to 8 ±2.8 h and 5 ± 0.58 h for siDLC3sp and siDLC3ss, respectively. This analysis shows that the increased RhoB levels observed upon DLC3 depletion are attributed to increased protein stability. RhoB was reported to be degraded via both the proteasomal and the endo-lysosomal pathways (Gong et al., 2018; Kovačević et al., 2018; Pérez-Sala et al., 2009). Indeed, treatment with MG132, a proteasomal inhibitor, or NH4Cl, an inhibitor of lysosomal acidification, increased RhoB levels in control cells (Figure 3 – figure supplement 3C). In Solo-depleted cells, RhoB levels were similarly increased by both drugs (Figure 3 – figure supplement 3C and 3D), indicating that both the proteasomal and the endo-lysosomal pathway are involved in RhoB protein degradation in cells with Solo downregulation.

In sum, these experiments show that Solo and DLC3 form a novel GEF – GAP pair that regulate endosomal RhoB in a complex manner, modulating not only its activity, but also localisation and turnover rates.

### Solo regulates EGFR trafficking and signalling

We previously found DLC3 to control EGFR degradation, a RTK that is rapidly endocytosed upon EGF ligand binding and is transported to the lysosomes in a RhoB-dependent manner (Braun et al., 2015; Gampel et al., 1999; Tomas et al., 2014). Considering the functional connection between Solo and DLC3, we explored whether Solo is also required for the regulation of EGFR trafficking. To this end, HeLa cells were stimulated with EGF and subjected to immunofluorescence staining with an antibody recognising the extracellular part of EGFR. This was followed by permeabilisation and incubation with an antibody binding to the intracellular domain of the receptor, enabling us to visualise surface and total EGFR pools (Figures 4A and Figure 4 – figure supplement 1). These experiments revealed a stabilisation of EGFR in the Solo knockdown cells. Because the receptor trafficking route determines the signalling response we next examined activation of the EGFR and downstream pathways by immunoblotting. Consistent with the immunofluorescence studies, Solo knockdown cells displayed increased basal EGFR levels and maintained elevated levels of the receptor for longer times post EGF stimulation (Figures 4B and 4C). Interestingly, in spite of EGFR phosphorylation (Figure 4D), downstream activation of the MAPK and PI3K pathways was dampened in Solo knockdown cells, as measured by AKT (Figure 4E) and ERK phosphorylation (Figure 4F), respectively. This indicates that EGFR cannot engage in efficient signalling under these conditions, presumably resulting from EGFR mislocalisation and the absence of respective downstream effector molecules.

In sum, our experiments identify Solo – DLC3 as a functional GEF – GAP pair that regulates RhoB and endocytic trafficking, impinging on cellular signalling responses triggered by EGF.

## DISCUSSION

Understanding how defined pairs of GEF and GAPs come together to control Rho GTPase signalling in a precise spatiotemporal manner is a long-standing challenge. In this study, we established an unbiased microscopy RNAi screen using the Golgi complex as a sensor and identified Solo as a RhoGEF protein that balances the RhoGAP activity of DLC3. We further discovered that the interplay between Solo and DLC3 is pivotal to the regulation of RhoB, a GTPase of central importance for endocytic trafficking.

Solo was previously described as a RhoA/C-targeting GEF involved in mechanotransduction, by binding and organising keratin-8/keratin-18 (K8/K18) filaments, thereby mediating force-induced RhoA activation and stress fibre formation (Abiko et al., 2015; Curtis et al., 2004; Fujiwara et al., 2016). Here, we uncover a previously unknown function of Solo in maintaining the integrity of the Golgi complex. Both the actin and microtubule cytoskeleton play a major role in the structure, function and positioning of the Golgi complex (Egea et al., 2013; Yadav and Linstedt, 2011). In addition, a few studies also suggest that an intact keratin network is required for the maintenance of Golgi structure (Fujiwara et al., 2016; Kumemura et al., 2004; Marceau et al., 2001). While our work did not focus on the organisation of cytokeratin filaments, we did find that Solo depletion reduces stress fibre formation in HeLa cells (Figure S1E), indicating that the Golgi phenotype we observe here may be a consequence of cytoskeletal rearrangements downstream of altered Rho activity. Importantly, out of the 23 RhoGEFs we tested in our imaging-based screen, Solo stood out for its ability to antagonise the fragmented Golgi phenotype caused by DLC3 depletion. This demonstrates that the effect of Golgi morphology is specifically downstream of the Solo – DLC3 interplay and not an unspecific readout of globally perturbed RhoGEF – RhoGAP balance.

The morphology of the Golgi complex integrates not only the organisation of the cytoskeleton but also membrane trafficking homeostasis. Among GTPases, in particular, RhoA, RhoB, RhoD, Rac and Cdc42 have been shown to affect various steps of membrane trafficking. We here find that endogenous Solo co-sediments in the membrane fraction together with RhoB but not RhoA and that the overexpressed protein physically interacts with GDP-locked RhoB. Interestingly, the GEF activity of Solo appeared to be important for the vesicular localisation of Solo since the GEF inactive mutant displayed a diffuse cytoplasmic distribution. This hints at a positive feedback loop between Solo activation and membrane enrichment. Such a mechanism was previously reported for the Sec7 Arf-GEF at the TGN (Richardson and Fromme, 2012). This GEF-dependent change in the vesicular localisation of Solo was also observed in HUVEC cells (Abiko et al., 2015), suggesting that the involvement of Solo in endocytic trafficking is conserved in other cellular systems as well. In line with this, Solo was contained in the endosomal mass spectrometry dataset of MEF cells obtained by Ivaska and co-workers (Alanko et al., 2015). While these studies do not allow to discriminate whether Solo is an endosomal cargo or an activator of RhoB, our optogenetic approach enabled us to address endosomal RhoB activation by Solo-GEF in a localised manner. To our knowledge, this is the first report of a direct readout of endogenous endosomal RhoB activation. Although we cannot exclude that Solo impinges on membrane trafficking by regulation of RhoA as well, our study clearly demonstrates a role of Solo in the regulation of RhoB. Since most of the endogenous Solo is membrane attached, for the future it will be interesting to explore the cellular cues that regulate the activation of full-length Solo on endomembranes.

We further uncover a multifactorial involvement of Solo and DLC3 in the regulation of RhoB, which extends beyond the regulation of its Rho GTPase activity. Specifically, we find that the interplay between Solo and DLC3 controls the activity, the subcellular distribution as well as the total protein levels of RhoB. Although we were not able to pinpoint the mechanistic details responsible for RhoB protein level regulation, an intact endosomal trafficking system was recently found to be required for the homeostasis of cellular RhoB amounts and protein localisation (Gong et al., 2018; Zaoui et al., 2019). This indicates that Solo and DLC3 may not only control RhoB activity directly but also indirectly by impinging on the endocytic recycling compartment.

Motivated by the interplay between Solo and RhoB, we also investigated the role of Solo in EGFR signalling and trafficking. Mirroring the effects of DLC3 depletion (Braun et al., 2015), Solo knockdown dampened AKT activation in response to EGF stimulation. These results indicate that the cellular levels of Solo play a critical role in determining the trafficking route of EGFR, hence the signalling potency of the receptor. The exact molecular mechanism behind this effect is, however, still open. Impairment of EGFR endocytosis using a dominant-negative dynamin mutant dampens PI3K/AKT and ERK signalling, leading to inhibition of EGF-dependent mitogenesis (Vieira et al., 1996). This is in line with our microscopy studies in HeLa cells where we observed delayed EGFR internalisation and signalling upon Solo knockdown.

The fact that both Solo and DLC3 knockdown delay EGFR trafficking while leading to opposing downstream signalling outcomes, illustrates the delicate GEF – GAP balance that is in place to control the trafficking and signalling capacity of this receptor. Besides the EGFR, it is plausible that also other cargoes are regulated by Solo and DLC3. For the future, it would be interesting to identify these through a more global approach. This strategy is expected to deliver insights into the cellular processes that are commonly regulated by Solo and DLC3, plus additional functions the two proteins might have that are independent of each other.

## ACKNOWLEDGEMENTS

We are thankful to Kai Hirzel and Angelika Hausser for help with setting up the optogenetics experiments. We are thankful to Angelika Hausser for helpful discussions and critical reading of the manuscript.

## AUTHOR CONTRIBUTIONS

Conceptualization: M.A.O., C.L.; Methodology: F.M., M.H., S.A.E.; Formal analysis: C.L., F.M.; Investigation: C.L., F.M., J.S., A.B., B.N., S.S., D.B., F.F.; Data curation: C.L.; Resources: S.S., S.A.E., M.H.; Writing original draft: C.L., M.A.O.; Writing - review & editing: C.L., M.A.O.; Visualization: C.L., F.M., M.H.; Supervision: M.A.O. Project administration: M.A.O.; Funding acquisition: M.A.O.

## MATERIALS AVAILABILITY

This study generated the following new plasmids: pIRESNEO-Solo L1217E-HA, pEGFP-RhoB T19N and CRY2-Solo-GEF-mCherry. Plasmids generated in this study as well as the anti-Solo antibody are available upon reasonable request. Further information and requests for plasmids, resources, and reagents should be directed to the lead contact Monilola A. Olayioye.

---

**Figure 1 – figure supplement 1:**
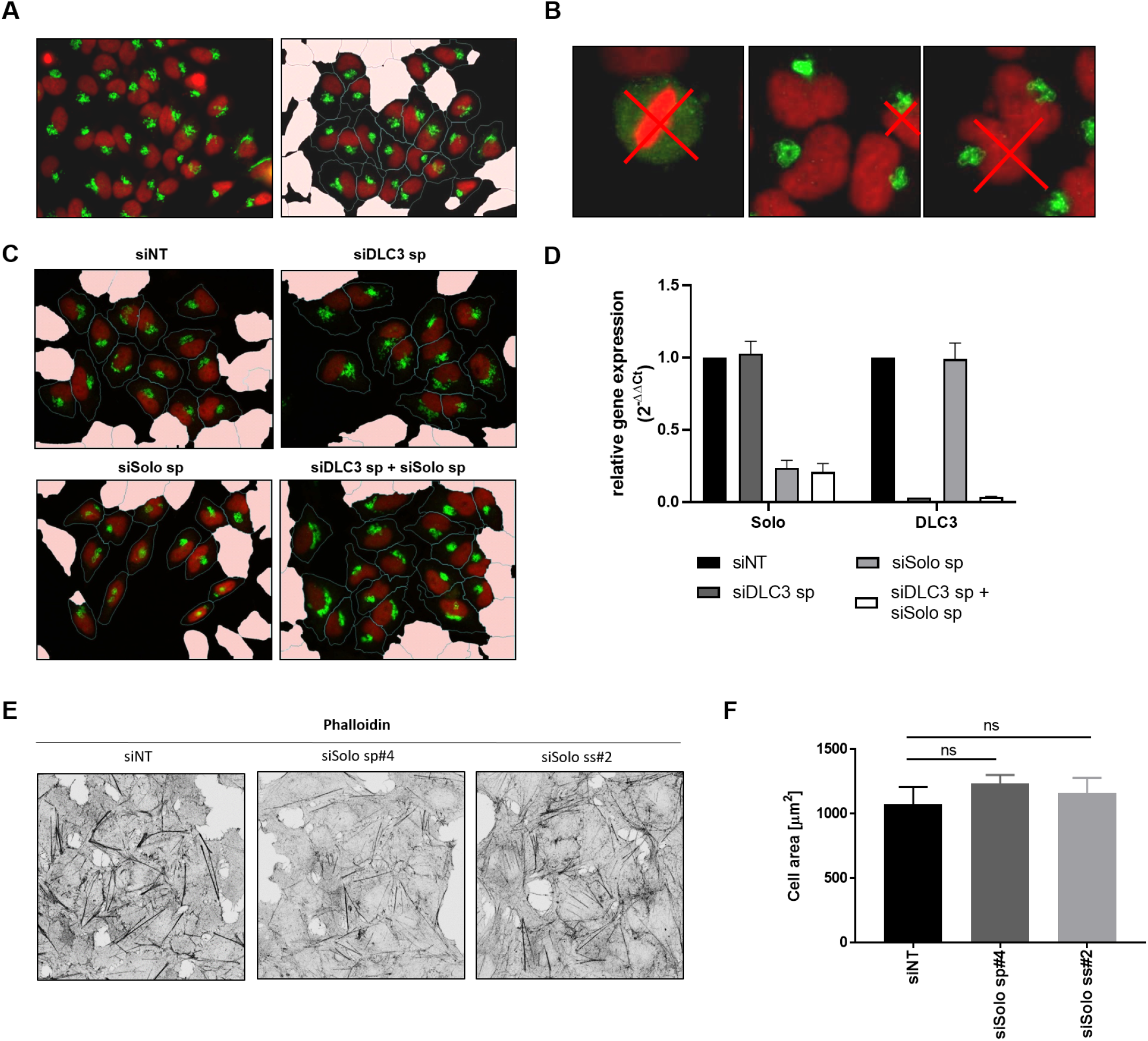
Imaging analysis pipeline for the automatic detection and characterization of Golgi structural parameters. (**A**) Representative fluorescence microscopy image acquired with the Hermes WiScan device pre- (left) and post- (right) processing via cell segmentation. HeLa cells were stained for the Golgi marker protein giantin (green). Nuclei were stained with TO-PRO-3 (red). Established criteria (see B) were used to define outlier cells, which are shaded in pink in the post-processed image. The dashed lines mark the outline of the cells. (**B**) Parameters used to classify cells as outliers. Mitotic cells (left panel), cells touching the image borders (middle panel) and cells that could not be segmented (right panel), as marked with the red crosses, were excluded from follow-up analysis. In addition, cells without detectable Golgi staining and cells with fragmented nuclei (not shown) were not taken into consideration. (**C**) Representative fluorescence microscopy images collected while performing the RhoGEF screen and demonstrating the successful implementation of the image segmentation algorithm established above. Shown are segmented HeLa cells fixed and stained for the Golgi complex (green) and nucleus (red) at 72 h post transient knockdown with the indicated siRNAs. (**D**) qRT-PCR measurements of DLC3 and Solo transcript levels in HeLa cells treated with the indicated siRNAs. The RNA was extracted at 72 h post transfection. Values were normalised to the reference gene RPLP0. n = 3, error bars: SEM. (**E**) HeLa cells were transiently transfected with the indicated siRNAs and incubated for 72 h before fixation and labelling of F-actin with Alexa Fluor488-phalloidin. Representative maximum intensity projections of 5 confocal sections are shown. All images were acquired and displayed using identical settings. (**F**) ImageJ quantification of cell area for images representatively shown in E. n = 82 – 118, N = 4. The significance of differences was analysed by a one-way ANOVA followed by a Dunnett multiple comparison test; n.s.: non-significant. For statistical testing data see Table S2. Refers to Figure 1.

**Figure 1 – figure supplement 2:**
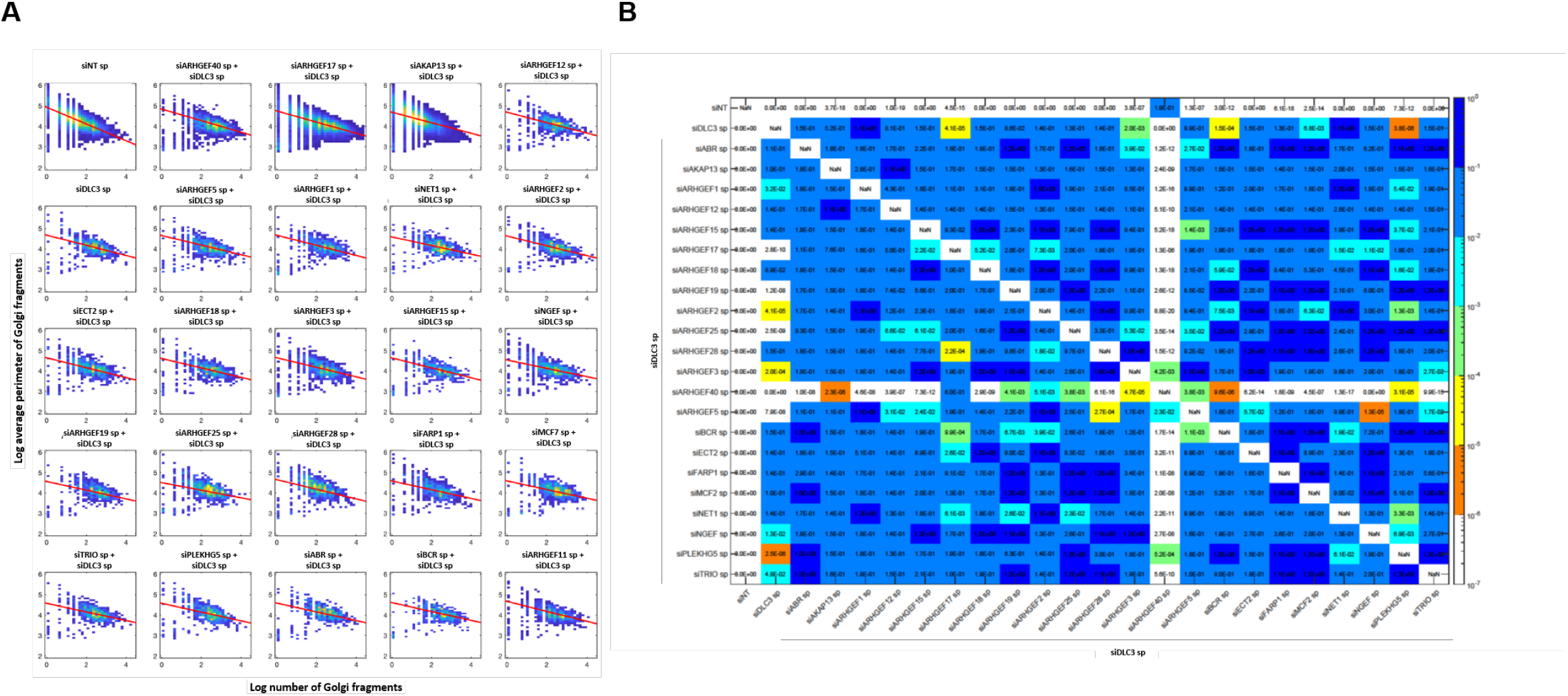
Data distribution and statistical testing of the Golgi imaging screen. (**A**) The normalised 2D histograms show the relation between the number (x-axis) and the average perimeter of Golgi vesicles (y-axis) of all cells observed, the control (siNT), the siDLC3, and the combination of DLC3 and RhoGEF specific siRNAs. Both axes are logarithmically scaled. The red lines indicated the least-square fits of the data. (**B**) Overview of the p-values obtained from the statistical analysis of the screen using NANOVA: The top-right values of the matrix shows the p-values of the average perimeter of Golgi vesicles, and the bottom-left values shows the p-values of the number of Golgi vesicles. The colour scheme indicates p-values logarithmically sorted. Refers to Figure 1.

**Figure 1 – figure supplement 3:**
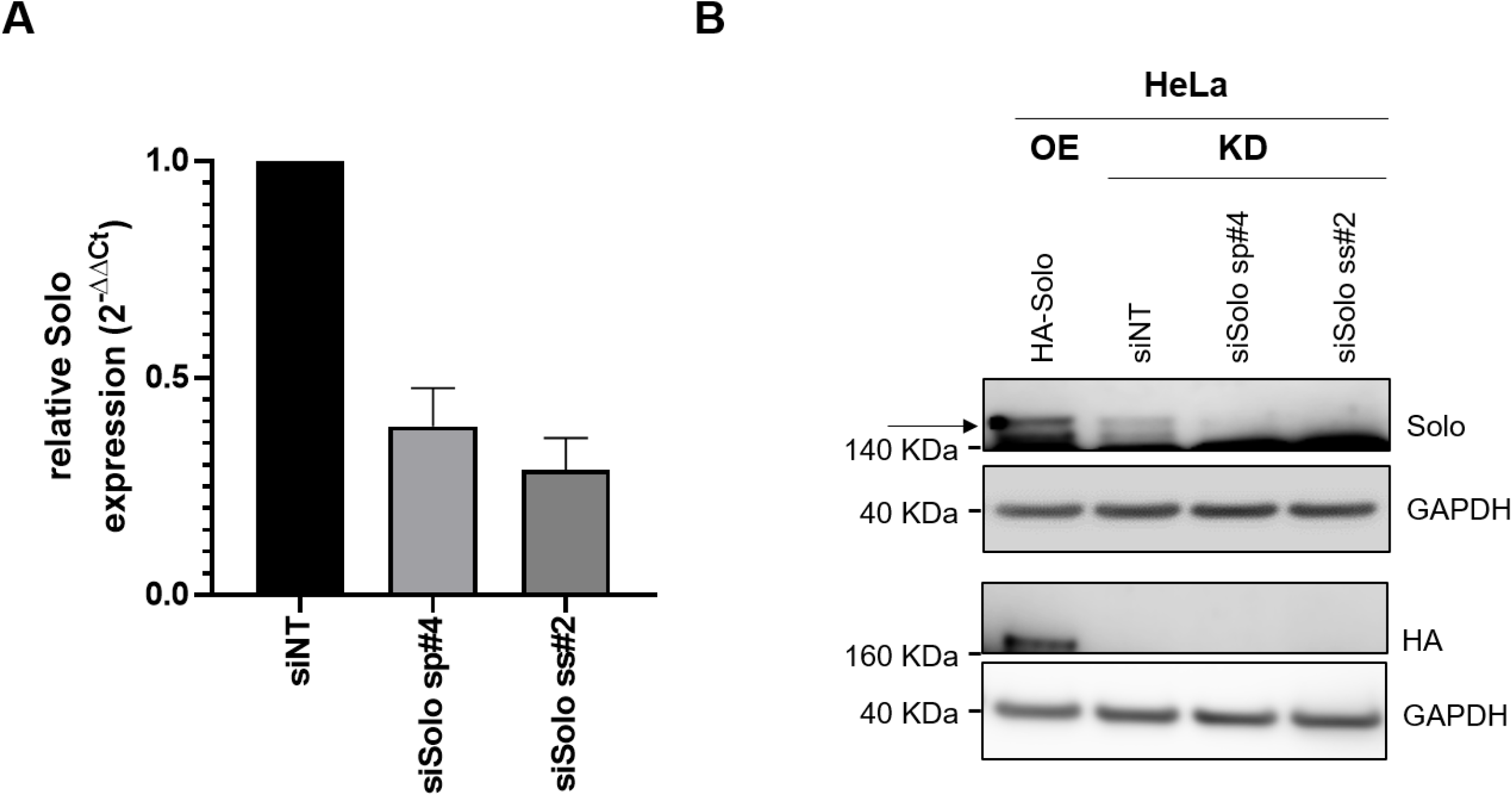
Validation of the Solo siRNAs and the Solo antibody. (**A**) qRT-PCR validation of two independent siRNAs targeting the Solo transcript. HeLa cells were transiently transfected with either control (siNT) or with Solo siRNAs (siSolo sp#4 and siSolo ss#2). The RNA was extracted at 72 h post transfection. The expression of Solo in the knockdown samples is plotted relative to the one in the sNT control cells, which was set to 1. Values were normalised to the housekeeping gene RPLP0. n = 3, error bars: SEM. (**B**) HeLa cells were transiently transfected with either a plasmid encoding for HA-Solo or with the indicated siRNAs. Full cell lysates were prepared at 24 h and 72 h post transfection, respectively, and probed by immunoblotting with the indicated antibodies. The band corresponding to Solo is annotated with an arrow. GAPDH was used as a loading control. Refers to Figure 1.

**Figure 2 – figure supplement 1:**
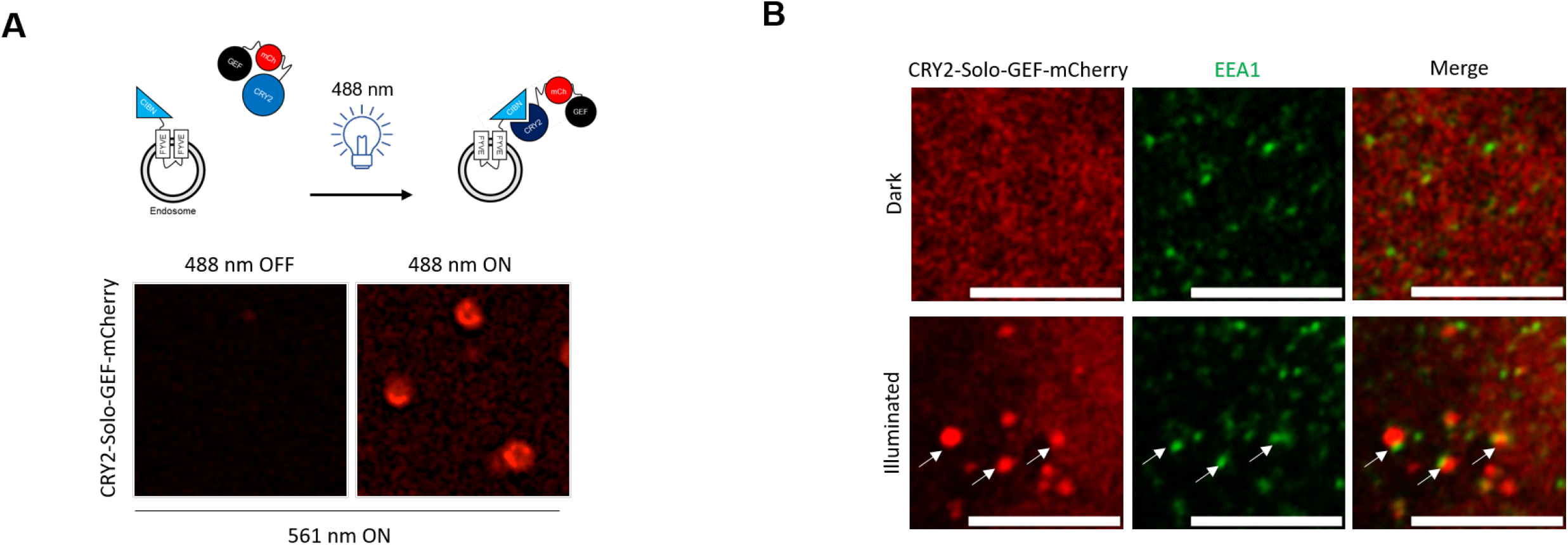
Validation of OptoEndo-Solo-GEF. (**A**) HeLa cells were transiently transfected with plasmids encoding CRY2-Solo-GEF-mCherry and CYBN-FYVE followed by imaging 6 h later. Shown are selected time frames from the live cell imaging experiments. mCherry was visualised by exposing the cells to 561 nm light. Before illumination with the 488 nm light, the CRY2-Solo-GEF-mCherry construct showed a diffuse and weakly fluorescent cytoplasmic localisation (left). Illumination triggered the acute accumulation of the probe to vesicles (right). The ON image was taken 48 sec after exposing the cells to 488 nm light. Images were acquired and are displayed using identical settings. (**B**) HeLa cells were transfected with plasmids encoding CRY2-Solo-GEF-mCherry and CYBN-FYVE followed by exposure to 405 nm light (5 sec ON – 35 sec OFF, 90 cycles) 14 h later, using a custom –made illumination box. A matching set of cells that were not exposed to light were prepared in parallel as dark control. Cells were fixed and stained for the early endosomal marker EEA1. White arrows show exemplary enrichment sites of the CRY2-Solo-GEF-mCherry probe on endosomes. Images were acquired and are displayed using identical settings. Scale bar: 5 μm. Refers to Figure 2.

**Figure 3 – figure supplement 1: Solo and DLC3 transiently co-localize in HeLa cells.** Live cell imaging of HeLa cells 24 h post transient transfection with plasmids expressing GFP-Solo (green) and mCherry-DLC3 K725E (red). Refers to Figure 3.

**Figure 3 – figure supplement 2:**
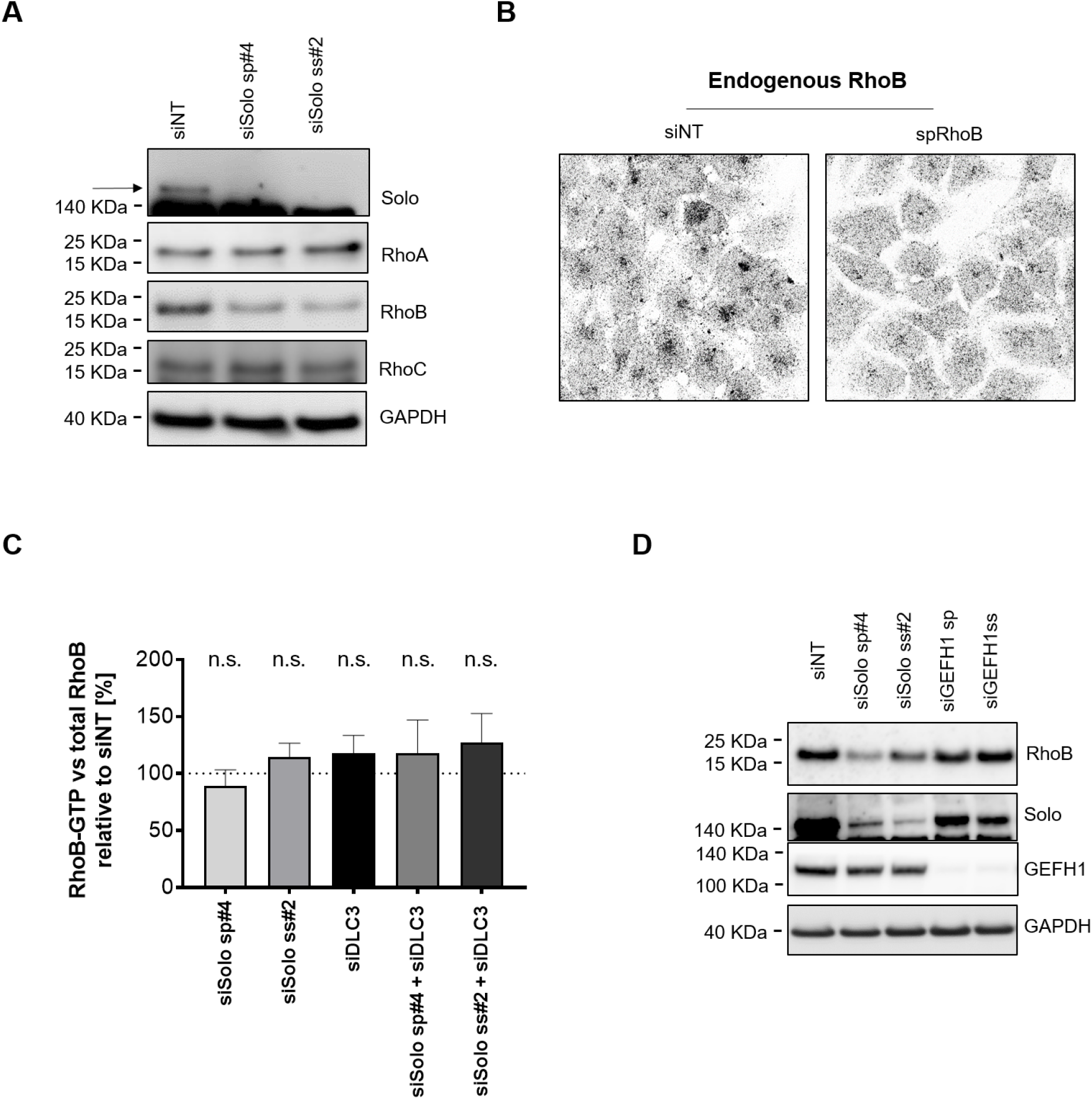
Solo and DLC3 co-regulate RhoB. (**A**) HeLa cells were transfected with the indicated siRNAs. At 72 h post knockdown, cell lysates were prepared and analysed by immunoblotting with the indicated antibodies. GAPDH was used as a loading control. (**B**) HeLa cells were transfected with either control (siNT) or RhoB (spRhoB) targeting siRNAs. 72 h post transfection, the cells were fixed and stained for RhoB. Shown are representative maximum intensity projections of 5 confocal sections. Both images were acquired and displayed using identical settings. (**C**) Further quantification of the RBD pulldown experiments shown in Figure 3. The RhoB-GTP pulldown signal was normalised to total RhoB and GAPDH and further divided by the signal obtained for the siNT sample. The value obtained for the siNT control was set as 100%, marked with the dotted line. The significance of differences was analysed by a one-way ANOVA and Dunett post-test; n.s.: non-significant. Control is the siNT sample. n = 3 – 5. (**D**) HeLa cells were transfected with the indicated siRNAs. At 72 h post knockdown, cell lysates were prepared and analysed by immunoblotting with the indicated antibodies. GAPDH was used as a loading control. For statistical testing data see Table S2. Refers to Figure 3.

**Figure 3 – figure supplement 3:**
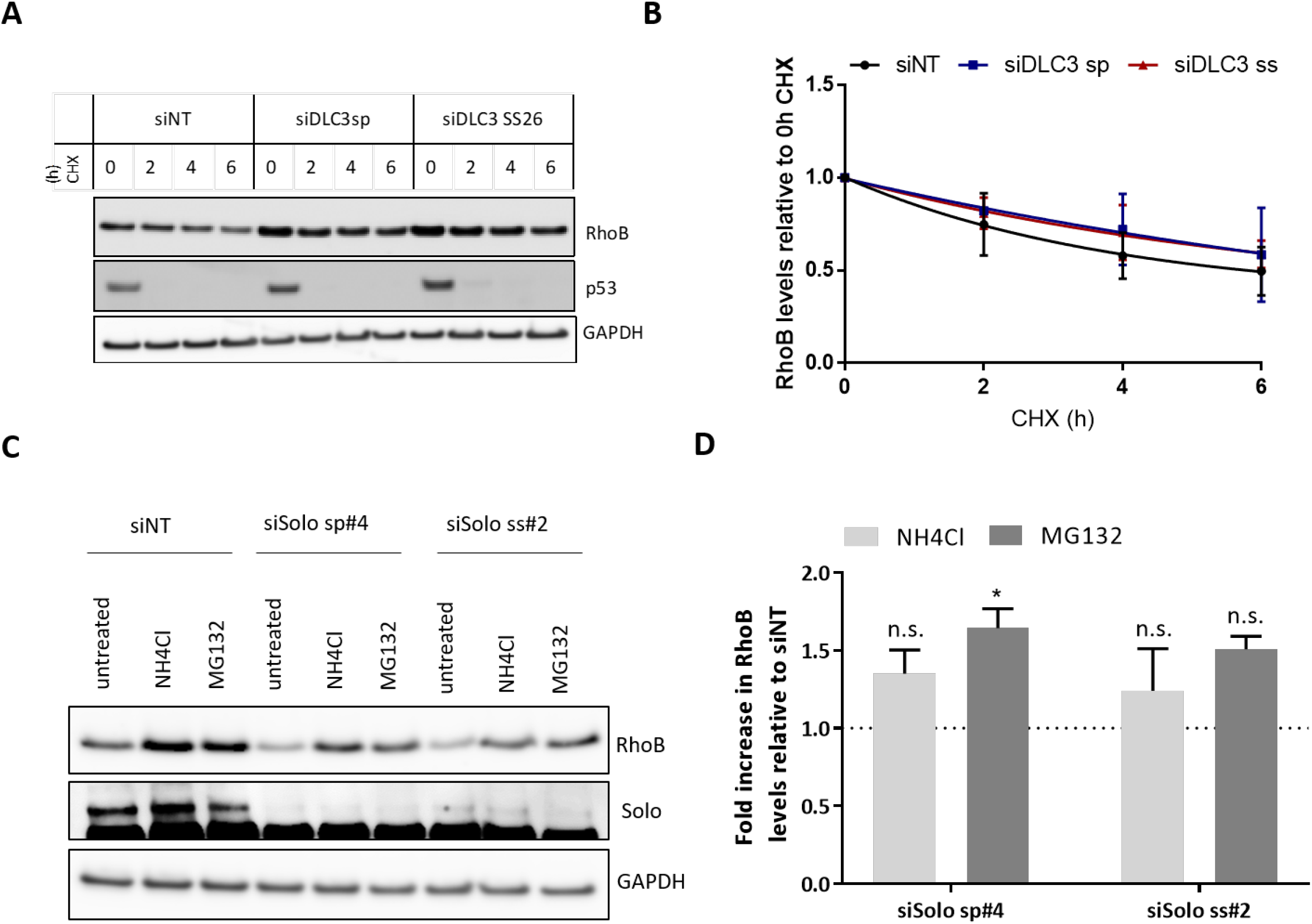
Solo and DLC3 regulate RhoB protein stability. (**A**) HeLa cells were transfected with the indicated siRNAs. At 48 h post knockdown, the cells were treated with 50 μg/mL cycloheximide followed by lysis at the indicated time points. P53 was used as a positive control for the treatment due to its short protein half-life. (**B**) Quantification of RhoB degradation kinetics for the cycloheximide treatment experiments representatively shown in A. Single exponential decay curves were determined using GraphPad Prism. n = 3 (**C**) HeLa cells were transfected with the indicated siRNAs. At 48 h post knockdown, the cells were treated with either 30 mM NH_4_Cl or 5 μM MG132. The cells were harvested 12 h later and the lysates were analysed by immunoblotting with the corresponding antibodies. (**D**) Densitometric quantification of RhoB levels of the experiments representatively shown C. Shown is the fold increase in RhoB levels in HeLa cells with Solo depletion vs. siNT control cells. The value obtained for the siNT control was set to 1, marked with the dotted line. Data of three independent experiments per condition were analysed by one-way ANOVA followed by Tukey’s post-test. *p≤ 0.03, n.s.: non-significant. Control is the siNT sample. n = 3. For statistical testing data see Table S2. Refers to Figure 3.

**Figure 4 – figure supplement 1:**
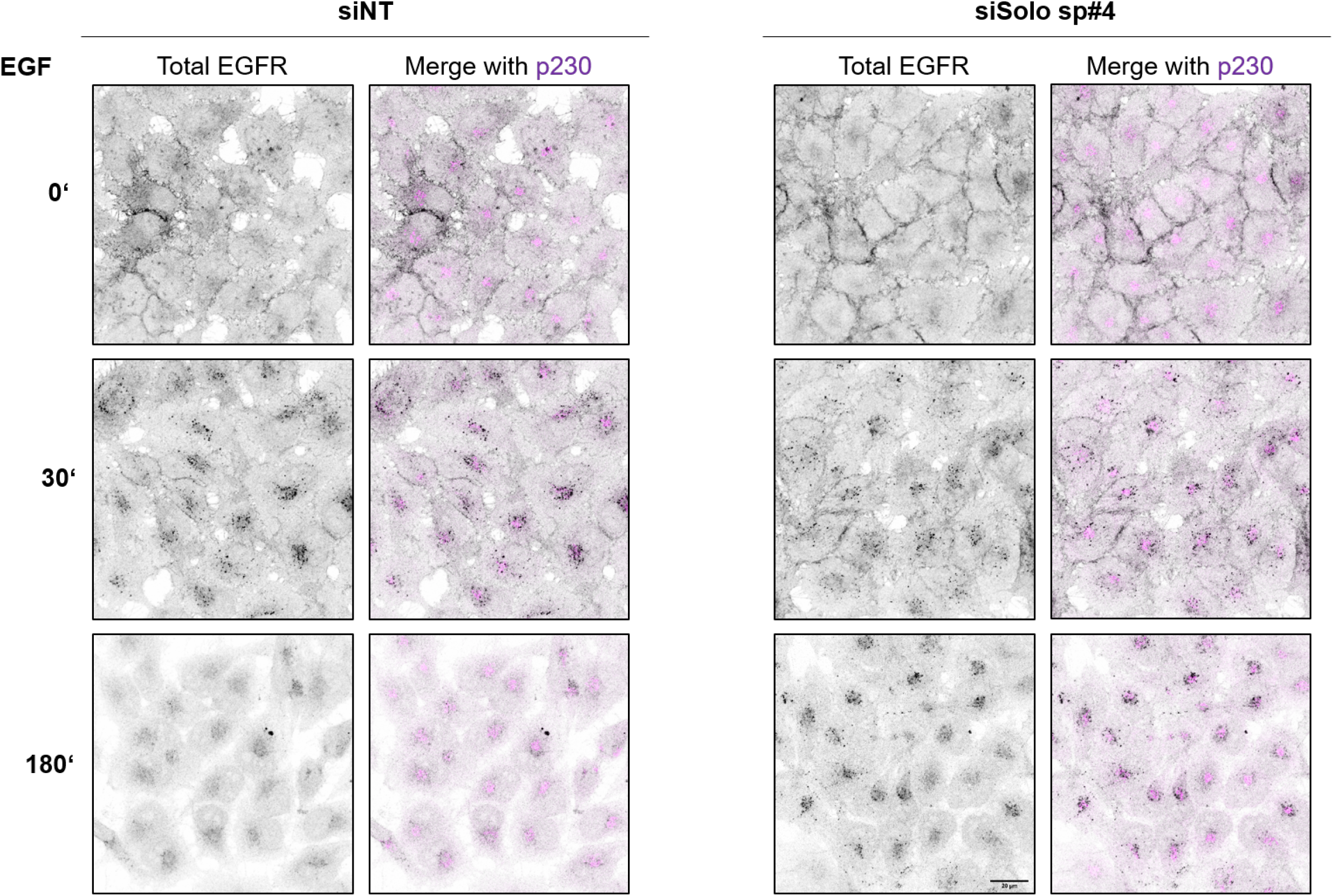
Solo knockdown delays EGFR trafficking. HeLa cells were transiently transfected with control (siNT) or Solo-specific (siSolo sp#4) siRNAs. Following a 48 h incubation period, the cells were serum starved overnight and, prior to fixation, stimulated with 10 ng/ml EGF for the indicated times. The cells were then stained with an antibody recognising the intracellular domain of EGFR (black) and the Golgi marker protein p230 (pink). Shown are representative maximum intensity projections of 5 confocal sections. All images were acquired and displayed using identical settings. Refers to Figure 4.

